# Ultra-precise quantification of mRNA targets across a broad dynamic range with nanoreactor beads

**DOI:** 10.1101/2020.11.05.369637

**Authors:** Ivan F. Loncarevic, Susanne Toepfer, Stephan Hubold, Susanne Klinger, Lea Kanitz, Thomas Ellinger, Katrin Steinmetzer, Thomas Ernst, Andreas Hochhaus, Eugen Ermantraut

## Abstract

Precise quantification of molecular targets in a biological sample across a wide dynamic range is a key requirement in many diagnostic procedures, such as monitoring response to therapy or detection of measurable residual disease. State of the art digital PCR assays provide for a dynamic range of four orders of magnitude. However digital assays are complex and require sophisticated microfluidic tools. Here we present an assay format that enables ultra-precise quantification of RNA targets in a single measurement across a dynamic range of more than seven orders of magnitude. The approach is based on hydrogel beads that provide for microfluidic free compartmentalization of the sample as they are used as digital nanoreactors for reverse transcription, PCR amplification and combined real time and digital detection of gene transcripts. We have applied these nanoreactors for establishing an assay for the detection and quantification of BCR-ABL1 fusion transcripts. The assay has been characterized for its accuracy and linear dynamic range. A comparison of the new method against the current clinical standard procedure with clinical samples from patients with chronic myeloid leukemia (CML) revealed good agreement between the results obtained with the respective method.

## Introduction

Digital amplification techniques [1] such a droplet digital PCR [2] or PCR on nanofluidic chips [3] offer the advantage of absolute quantification with exquisite precision [4]. Precision, and as such the measurement range of digital assays is a direct function of the number of partitions used for the analysis [5]. The more extensive the dynamic range the more partitions must be used to meet the Poisson criteria for analysis. However microfluidic tools are needed to split a sample into sub-nanoliter droplets or micro structured substrates for distributing the sample across nano-chamber arrays. Endpoint fluorescence generated by the amplification reaction in each respective partition is used to detect partitions containing amplification targets. Recently, we [6] and others [7, 8] have introduced techniques for generating digital compartments with hydrogel beads serving as volume templates for aqueous partitions of the sample and thus allowing for microfluidics-free digital assays. A schematic of the developed workflow is shown in Figure 1.

**Fig 1.**
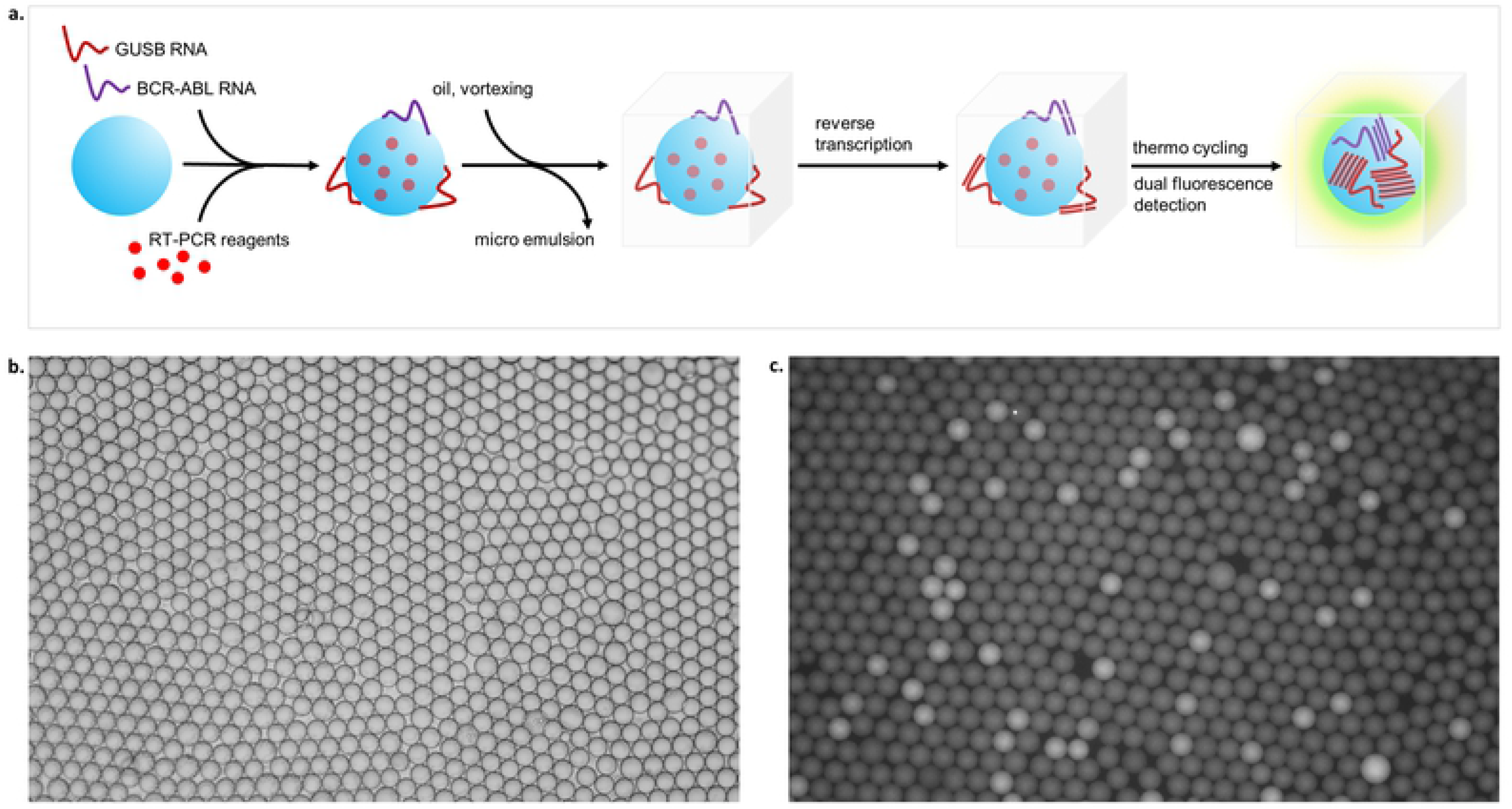
Bead Nanoreactor Workflow. a.) Process steps: i.) bead nanoreactor loading with a solution containing amplification reagents and the extracted nucleic acid. ii.) bead transfer to non-aqueous liquid with a suitable emulsifier. iii.) cDNA synthesis on encapsulated hydrogel bead. iv.) bead melting, PCR and fluorescence detection. b.) bright field image of bead nanoreactors in PicoSurf oil. c.) fluorescence image of nanoreactor beads post PCR

In this work we are employing monodisperse non-crosslinked hydrogel beads comprised of agarose and gelatin. The composition has been found mechanically stable to sustain the handling during the assay procedure and to provide for RNA template binding in RT-PCR buffer. Nanoreactor beads provided in RT-PCR buffer are exposed to extracted RNA mixed with RT-PCR reagents. The hydrogel composition is designed to allow the reagents to diffuse into the interior of the beads. Following an incubation step for reagent take-up and nucleic acid binding the beads are transferred to a fluorocarbon oil containing an emulsifier by extensive shaking. The bead suspension is then transferred to a detection chamber designed (shown in Figure 2) to conveniently generate a single bead layer for detection of the fluorescence within the individual partitions created by the beads. The detection chamber features a luer-lock inlet and a luer-lock outlet on a frame with an attached polycarbonate sheet with a thickness of 100μm and transparent cover part on the opposite side of the frame. The assembly forms a flat chamber between the polycarbonate sheet and the transparent cover with a height of approximately 120μm. The detection chamber is filled with the bead emulsion and clamped on a temperature control unit whereby the thin polycarbonate layer is brought into close contact with the surface of a Peltier element. The Peltier element is part of a custom made temperature control unit that is designed to fit on a standard microtiter plate interface on an x,y-stage of an inverted fluorescent microscope equipped with fluorescent filter sets for FAM, Cy3 and Cy5 dyes and with a CMOS camera for fluorescence imaging.

**Fig 2.**
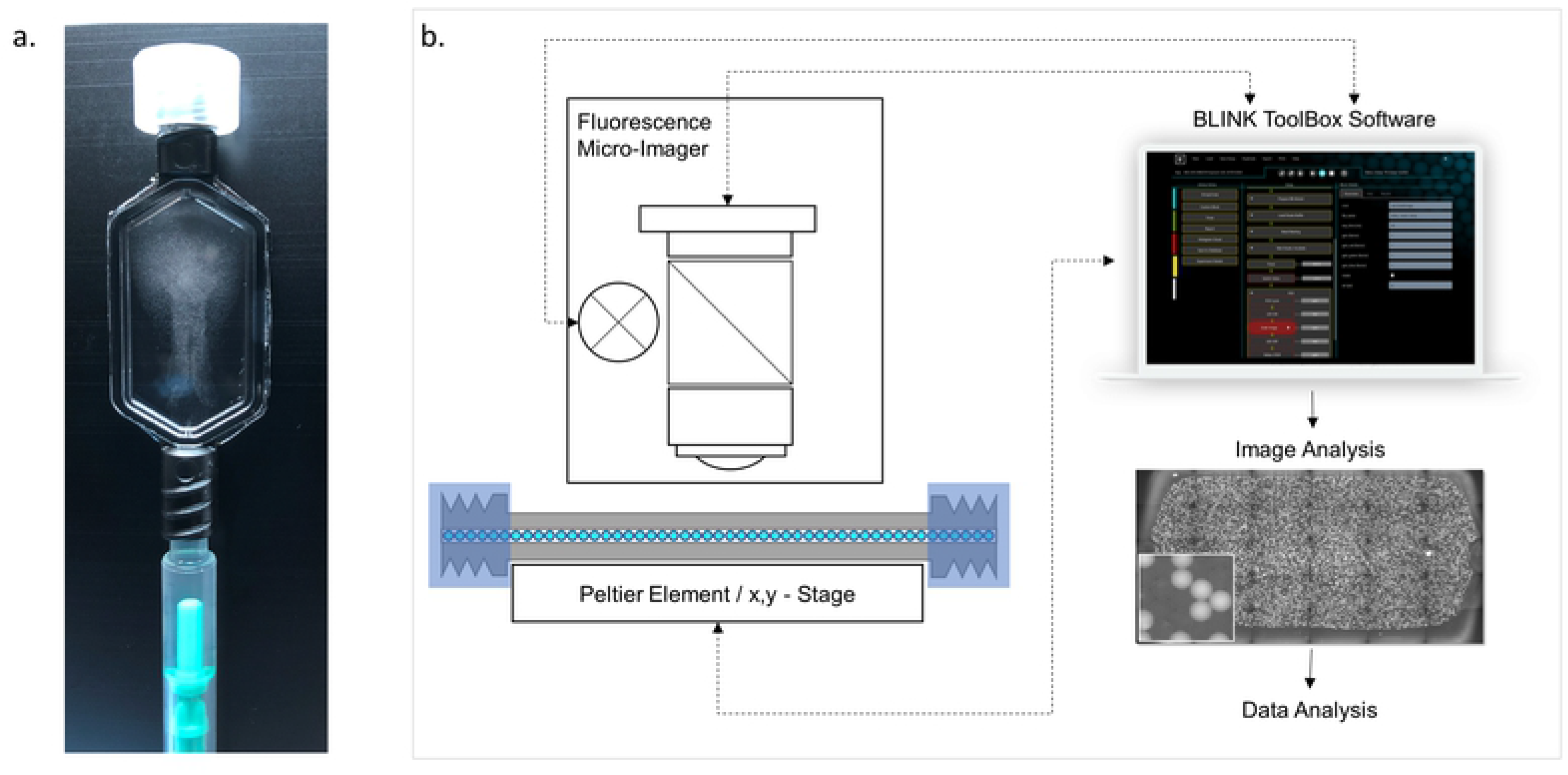
Incubation and detection set-up. a.) Incubation chamber, in the process of bead loading with a syringe; the bead suspension is visible in the reaction chamber. b.) schematic of the detection set-up comprising the incubation chamber mounted on a Peltier-element for thermocycling placed on a x,y-stage under an epi-fluorescence microscope; the process is automated through the Blink ToolBox Software package

The set-up has been applied to establish a detection format that combines endpoint (digital) and real time detection in each individual nanoreactor compartment. We found that by analyzing only a modest number of beads if compared to other digital assay methods a highly precise assay with an unprecedented dynamic range can be realized [5, 9–11]. The method was applied to develop an assay for the quantification of the amount of BCR-ABL1 mRNA relative to the internal reference gene transcript GUSB [12]. Chronic myeloid leukemia (CML) is a myeloproliferative neoplasm characterized by the presence of the BCR-ABL1 fusion gene [13]. Quantification of the BCR-ABL1 transcript level reflects leukemic burden. Molecular response to treatment is determined based on the ratios of BCR-ABL1 and control gene transcript levels. Here we present the results of the evaluation of the new assay with nanoreactor beads and compare the data with results obtained with the current clinical gold standard.

## Materials and methods

### Nanoreactor beads

A solution containing 1.5% Acetone-insoluble gelatin from bovine skin type A G1890 (Sigma) or porcine skin type B G9391 (Sigma) and 0.5% Low-gelling 2-Hydroxyethyl agarose (A4018, Sigma) has been prepared in nuclease-free water (Carl Roth) and incubated at 50°C under gentle agitation (750rpm). The solution has been used to generate low dispersity microbeads on the μEncapsulator system (Dolomite microfluidics) using a fluorophobic droplet junction chip (100μm) with a 4-way linear connector to interface the fluidic connection between tubing and chip. Two Mitos P-Pumps were used to deliver the hydrogel solution and the carrier oil (Picosurf-1, Spherefluidics). The system was modified with an integrated heating rig which is placed on top of a hot plate allowing maintaining the gelatin/agarose hybrid solution in liquid state and heating up the driving fluid ensuring consistent temperature for all components and liquids. Picosurf-1 and the hybrid hydrogel solution were both pre-filtered with a 0.22μm filter before they were placed into their respective reservoirs. Temperature of the heating rig is set to 55°C to heat the gelatin-agarose hydrogel as well as the droplet junction chip. The nanoreactor beads were collected in a tube on ice and incubated for a minimum of 48h at 4-8°C before phase conversion. Thereafter, the excess of Picosurf-1 was removed from the nanoreactor bead emulsion and 1H,1H,2H,2H-perfluorooctanol (PFO; Sigma) was added to the same volume. The tube was vortexed to break the emulsion and centrifuged for 5s at 2,500 x g. The aqueous phase with the beads was transferred to a fresh tube and the PFO-procedure was repeated. Excess of PFO was removed by transferring again the aqueous phase with the beads into a new tube. The nanoreactor beads were washed 4x with an equal volume of nuclease-free water, aliquoted and stored at 4°-8°C for further use.

### Samples

Enrollment of patients in this study was approved by the Jena University Hospital (JUH) ethics committee (2719-12/09). RNA was extracted from peripheral blood and from cell lines at the molecular-oncology laboratory at Jena Unversity Hospital (JUH) using the TRIzol standard procedure [14, 15]. The total RNA concentration of the patient samples varied between 62-262 ng/μl. Samples with no, medium or high levels of typical e13a2 or e14a2 BCR-ABL1 transcripts were selected for a comparison analysis (Table 1 in supporting informantion). Each RNA stock was divided in two equal parts representing a reference sample and a sample to be processed on the newly developed nanoreactor beads. The reference sample was processed at the molecular-oncology laboratory at JUH. RT-PCR was performed as a two-step real time rtPCR with SuperScript™ IV VILO™ from ThermoFisher, followed by a PCR as described elsewhere ([15]. The Nanoreactor-sample was subjected to a one-step RT-PCR process. RNA from cell line sample HL60 (BCR-ABL1 negative) and K562 (e14a2 BCR-ABL1 positive) RNA with approximately 700ng/μl were mixed to obtain the desired ratios BCR-ABL1/GUSB in contrived samples for the precision analysis.

**Table 1.** Precision results for BCR-ABL1 and GUSB.

| Level | Replicates | Target | Mean [log10 cp/bead] | Mean* [cp/reaction] | SD [log10 cp/bead] | Analysis Method |
| --- | --- | --- | --- | --- | --- | --- |
| Low | 6 | BCR-ABL1 | -2.72 | 19 | 0.09 | Poisson |
|  |  | GUSB | 2.04 | 1,100,000 | 0.02 | Ct |
| Medium | 6 | BCR-ABL1 | -1.03 | 933 | 0.04 | Poisson |
|  |  | GUSB | 2.00 | 1,000,000 | 0.04 | Ct |
| High | 6 | BCR-ABL1 | 1.85 | 708,000 | 0.04 | Ct |
|  |  | GUSB | 2.10 | 1,260,000 | 0.02 | Ct |

\* for 10,000 beads per reaction, SD= standard deviation

**Table 2.** Precision results for ratios BCR-ABL1/GUSB.

| Level | Ratio BCR-ABL1/GUSB |  |  |  | Analysis Method |
| --- | --- | --- | --- | --- | --- |
|  | Mean [log10%] | Mean [%] | Mean MR | SD [log10%] |  |
| Low | -2.76 | 0.0017 | 4.76 | 0.09 | Poisson/Ct |
| Medium | -1.04 | 0.091 | 3.04 | 0.01 | Poisson/Ct |
| High | 1.75 | 56 | 0.25 | 0.06 | Ct/Ct |

(SD = standard deviation, MR = molecular response)

### Probes and primers

Primers and probes, each at 0.4μM, were used as described elsewhere (ß-Glucoronidase (GUSB), ABL1 [16], and BCR [17]). No additional oligonucleotides were used for reverse transcription in the nanoreactors.

### RT-PCR on nanoreactor beads

Nanoreactor beads were loaded with a solution containing target RNA and the RT-PCR reagents by mixing 40μl of sedimented beads with 10μl RNA sample and 50μl 2x RT-PCR solution for 8 min at 20°C at 1000rpm (100μl final volume). The concentration of the resulting 1x RT-PCR Mix was 20mM Tris HCl, 22mM KCl, 22mM NH4Cl, 3mM MgCl2, 0.4 U/μl Hot Start Taq DNA Polymerase (biotechrabbit GmbH), 0.4 mM dNTPs (biotechrabbit GmbH), 1x RT-Mix (biotechrabbit GmbH), 0.1% (w/v) BSA (Sigma), and 0.4μM TaqMan probes and primers for both GUSB and BCR-ABL1 amplifications. Thereafter beads were sedimented at 300g for 30 seconds and the supernatant was removed. Picosurf-1 (100μl) was added to the sediment (30-40μl) and an emulsion was produced by simply sliding the tube over the holes of a microcentrifuge tube rack 20 times at a frequency of approximately 20/s while pressing the tube against the rack surface or with 2x 5 seconds shaking at speed level 3 in the Minilys Homogenizer (Bertin Technologies). Microemulsion resulting from the excess of aqueous liquid not accumulated on the beads and forming a layer below the beads was removed. The latter was repeated after resuspending the beads and the microemulsion in another 100μl Picosurf-1. The bead suspension was transferred to the detection chamber that is placed on a PELTIER element (Quick-Ohm, Küpper & Co. GmbH, #QC-71-1.4-3.7M) in a customized test rig. The test rig is mounted on a motorized x,y-stage of an Axio Observer Microscope (Zeiss). Using the ToolBox Software (https://www.blink-dx.com/technology/toolbox) the parameters for thermocycling and imaging were programmed and the microscope, Peltier-Element and x,y-stage controlled. Reverse transcription was conducted for 10min at 50°C, followed by PCR (3min 95°C for initial denaturation and 40 cycles of 5sec 95°C, 5sec 60°C for amplification). Images were taken using a 5x objective (field of view 416mm x 2.774mm), pE-4000 (CoolLED Ltd.) light source and a CMOS camera (UI-3260CP-M-GL, IDS). Fluorochrome specific filters for FAM, Cy5 and Rox or Att550 F36-501, F36-523 and F36-560 (Semrock) were used for imaging in the respective fluorescence channel. Image acquisition was performed after each PCR cycle at 60°C at one discrete position of the detection chamber. At the end of the PCR run, the entire detection chamber was scanned at 20°C resulting in 40 individual image positions (end point; digital analysis). Figure 3 shows exemplary fluorescence images of the GUSB and BCR-ABL1 RNA amplification after one, twenty-nine and forty PCR cycles. The images derive from the precision analysis and a sample with 800 copies BCR-ABL1 and 900,000 copies GUSB per reaction (medium level as described in Results). Positive Beads approaching saturation are clearly visible for GUSB after 29 cycles, whereas beads containing single BCR-ABL1 targets show up weakly positive after 29 cycles.

**Fig 3.**
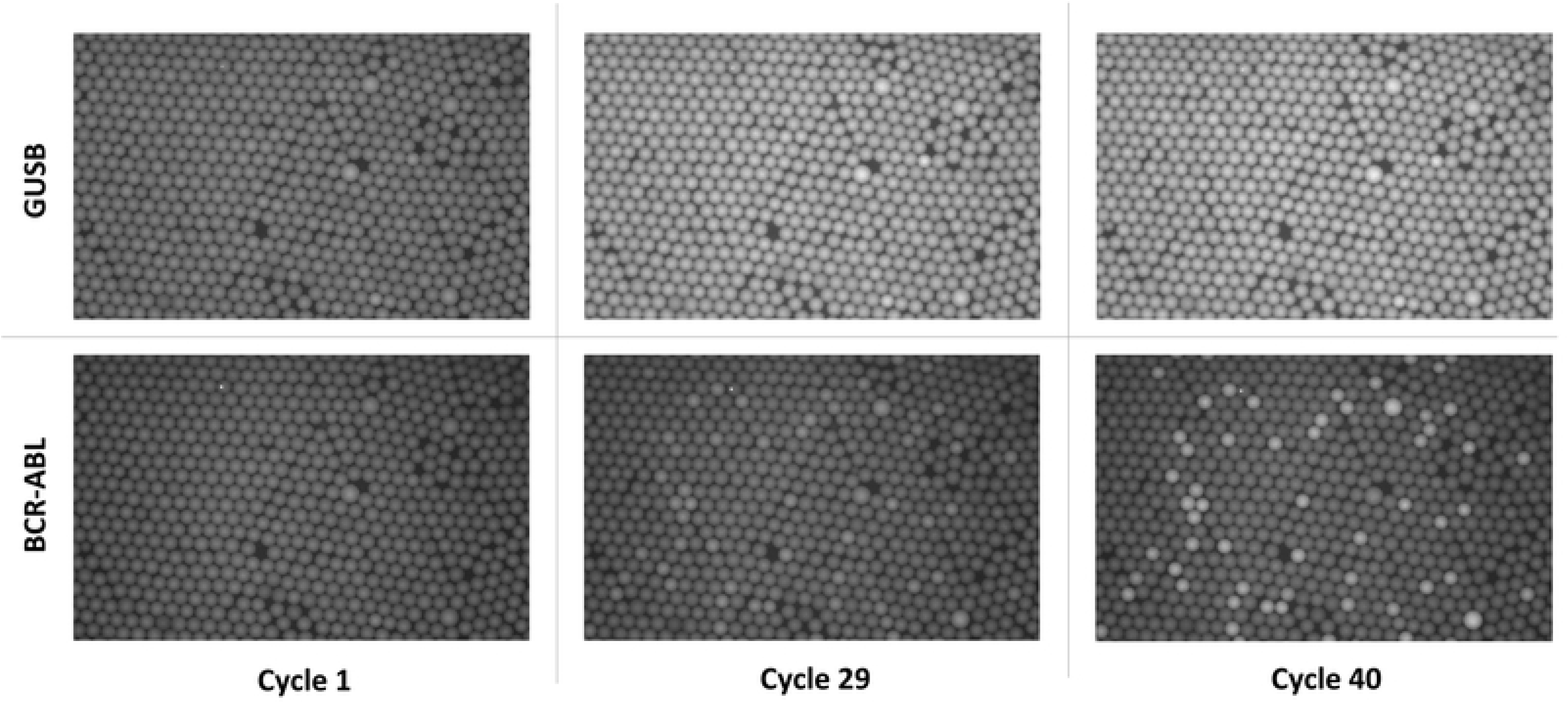
Fluorescence Images from real time PCR analysis. Fluorescence images after one, twenty-nine and forty cycles for the GUSB and BCR-ABL1 specific channels (FAM for GUSB, upper row; Cy5 for BCR-ABL1, lower row).

### Image data analysis

A segmentation algorithm [18] is used to identify the bright disk-shaped nanoreactor beads against dark background in fluorescence images. Circles are fitted to the identified bead contours. Grey value representative for signal intensity and size are determined for each bead. Each bead position is tracked in consecutive images for individual real time PCR analysis.

Target numbers are quantified using either digital PCR analysis based on Poisson statistics or on real-time quantitative PCR analysis by comparison against a calibration curve. The appropriate analysis method is determined by the number of positive beads in relation to the total number of beads identified in the detection chamber. For digital PCR, a fluorescence signal threshold is set to distinguish positive beads from negative beads. For real-time quantitative PCR, cycle threshold values are derived from a nonlinear model fitted to the time course of the fluorescence signal for each individual bead. The nonlinear model combines a sigmoid and a linear function where the sigmoid component reveals amplification kinetics, and the linear component represents baseline of the signal. The mean cycle threshold of all beads is a measure for target present in the sample. As a rule, we are applying Poisson analysis if the proportion of beads remaining negative after amplification is larger than or equal to 0.5%. Based on a Poisson distribution of targets across beads, a ratio of 0.5% negative beads corresponds to an average of 5.3 copies per bead. The resulting upper limit of the digital measuring range is 5.3*N, where N is the total number of beads. If the proportion of dark beads is smaller than this limit of 0.5%, real-time quantitative PCR is utilized.

Digital analysis is based on a Poisson correction to account for the fact that positive beads can contain more than one target. Bead volume variations are factored into the quantitation by performing bead volume specific Poisson corrections. Quantitation for each target is expressed in copies per bead. Real-time quantitative analysis calculates the target number per bead based on the mean cycle threshold value using analyte specific calibration parameters. The overall analysis procedure is summarized in Figure 4.

**Fig 4.**
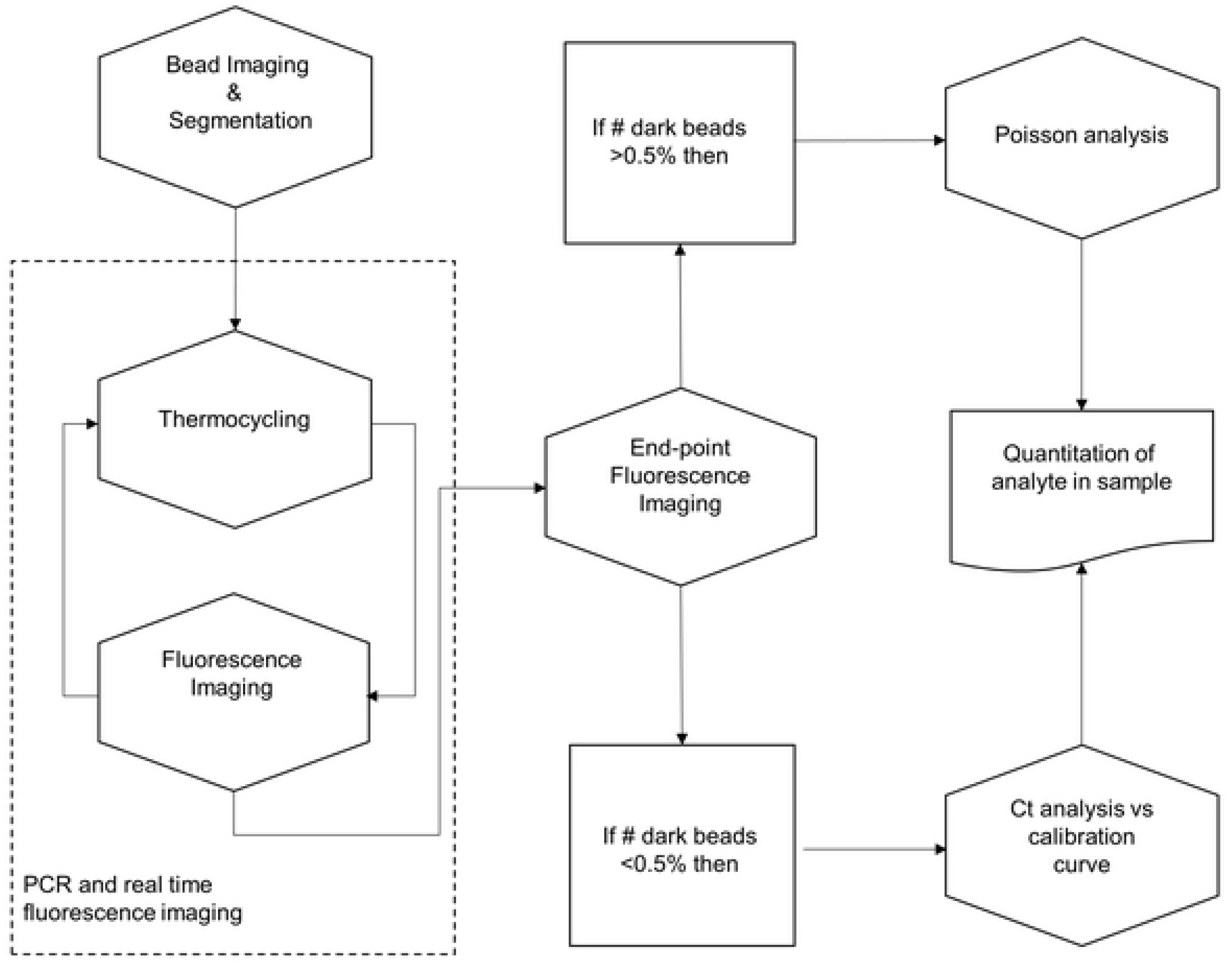
Analysis algorithm for combined digital and real time analysis.

## Results and discussion

We sought to establish an assay for the quantification of BCR-ABL1 and GUSB transcripts employing nanoreactor beads and characterized the assay for its precision and dynamic range. We also performed a method comparison against the current clinical standard with clinical samples from patients with CML [19]. The BCR-ABL1 and GUSB working standards were quantified based on droplets and nanoreactor beads and are traceable to the calibrant ERM-AD623 (Merck), a certified reference material for quantification of BCR-ABL1 and GUSB DNA (data not shown). The data analysis algorithm requires calibration curves for the quantification of high copy numbers via real-time quantitative PCR analysis. For this purpose, RNA dilution series with medium and high BCR-ABL1 and GUSB copy numbers were independently tested. The concentration levels represent the expected measuring range for clinal samples. The number of copies per bead ranged from 4 cp/bead to 118 cp/bead for BCR-ABL1 and from 7 cp/bead to 251 cp/bead for GUSB. For the estimation of calibration curve parameters, linear regression was performed to establish the correlation between median Ct values from all beads and input copies per bead on a logarithmic scale with formula *ct* = *offset* + *slope* · log10 *cp*/*bead*. The efficiencies for BCR-ABL1 and GUSB standard curves are 106.5% (slope −3.176) and 100.6% (slope −3.307), respectively. Offsets are 23.459 for BCR-ABL1 and 24.326 for GUSB. Coefficients of determination R^2^ are 0.9596 for BCR-ABL1 and 0.9761 for GUSB.

In order to assess the measurement range of BCR-ABL1 we tested the assay with 33 different contrived samples with a fixed GUSB concentration and titrated BCR-ABL1 concentration ranging from −3.44 log10 cp/bead (0.00036 cp/bead) to 2.83 log10 cp/bead (676 cp/bead). Quantification results are shown in Figure 5.

**Fig 5.**
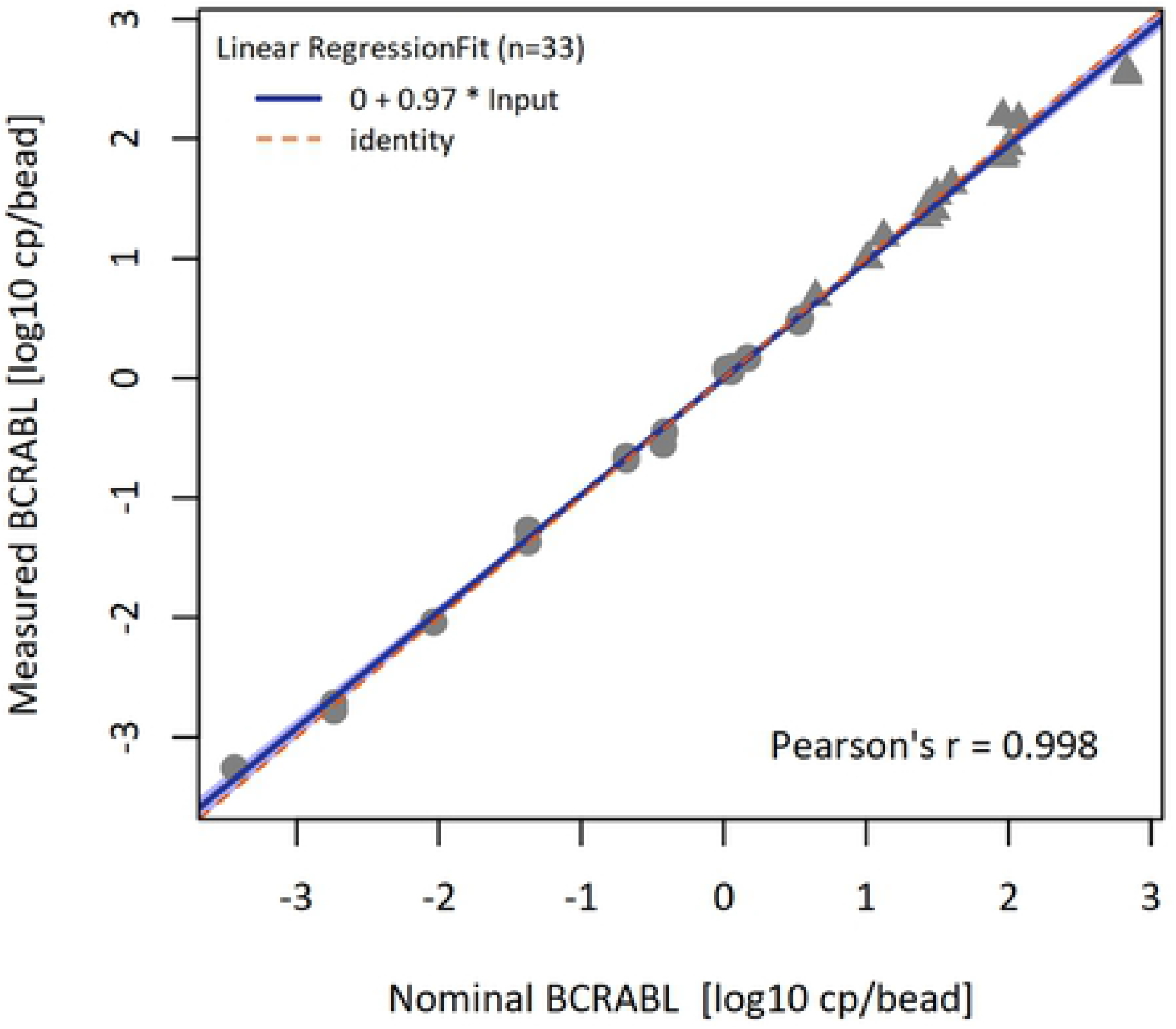
BCR-ABL1 measuring range. Input copies per bead are plotted on the x-axis against the measured copies per bead on the y-axis (both logarithmic scale; dots: Poisson analysis of digital PCR data, triangles: Ct-value analysis of quantitative real-time PCR)

The slope for the least-squares regression was determined at 0.97 with 95% confidence interval from 0.95 to 1.00. The estimated intercept was 0.00 with 95% CI from −0.03 to 0.03. R^2^ was 0.9965. As symbols indicate, measured results were calculated using either Poisson analysis (circles) from end point fluorescence imaging or Ct-value analysis from real-time fluorescence imaging (triangles). The data shows linearity and a high degree of concordance between input and measured BCR-ABL1 values across a measuring range of more than six orders of magnitude. For the given data set, real-time quantitative PCR extends the measuring range of digital PCR by more than two orders of magnitude.

We assessed the repeatability of the assay with three contrived samples comprising different ratios BCR-ABL1/GUSB (low, medium, high). The chosen BCR-ABL1/GUSB levels reflect the clinically relevant range to monitor the disease status in the context of CML therapy. Each sample was analyzed with six replicates. Replicates were tested in different laboratory runs, as the nanoreactor beads were configured to process one sample at a time. Therefore, repeatability conditions include between-run imprecision. Results for the precision of quantification for BCR-ABL1 (absolute) and GUSB (absolute) are summarized in Table 1. **Error! Reference source not found.** shows precision results for ratios BCR-ABL1/GUSB.

We used the following formulas to report ratios BCR-ABL1/GUSB:

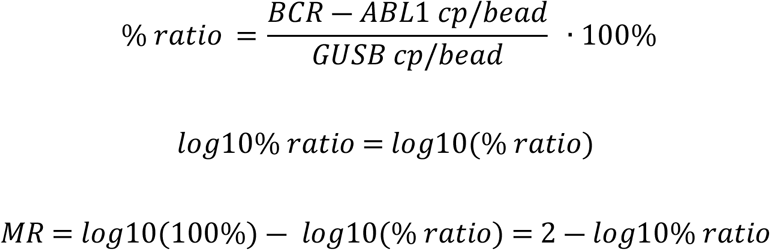

The molecular response (MR) is defined as the log reduction level of BCR-ABL1 compared to a control gene under consideration of the conversion factor to ensure traceability to an International Standard (IS). The molecular response (MR) was calculated without using the international scale, because a laboratory-specific conversion factor has not yet been established for the new method. Precision is expressed as standard deviation (SD) for each BCR-ABL1/GUSB level separately.

The concentration of GUSB target was set to a mean value of 2.05 log10 cp/bead for all levels resulting in approximately 1,100,000 GUSB copies in 10,000 beads. The lowest measured ratio BCR-ABL1 −2.76 log10% corresponds to a molecular response (MR) between 4.5 and 5. MR^4.5^ and MR^5^ is considered state of the art for the detection and quantification limit of the method for clinical applications. To ensure assay sensitivity required to achieve MR^5^, a minimum number of 240,000 copies of control gene transcript GUSB is recommended [20]. Because of the high capacity of the beads our assay is exceeding this recommendation by a factor of four.

In comparison, the current technically-leading commercially available test features a precision of 0.25 log10 for MR^≤4.6^ (QXDx™ BCR-ABL1 %IS Kit by Biorad digital PCR). This precision claim takes all variance components into account, including different instruments, reagent lots, operators, and analytical repeatability. However, repeatability is by far the strongest contributor to variance. Therefore, the achieved repeatability with the nanoreactor beads of 0.09 log10% for MR between 4.5 and 5 can be considered excellent. The implementation of real-time quantitative PCR analysis for nanoreactor beads provides favorable precision results because variance from the signal detection process is strongly reduced by averaging Ct-values from approximately 550 independent nanoreactors.

Peripheral blood samples from CML patients were used to compare test results obtained with the new assay format against standard laboratory results. All tests were performed with approximately 10,000 beads per run. With an allowed maximum of 5.3 targets per bead, the upper limit of quantification for the digital readout is 53.000 targets per test. For quantification of higher target numbers real-time quantitative PCR analysis based on the established calibration curve has been applied. A total of 28 clinical specimens from CML patients were provided by the Department of Hematology at JUH in Germany. The ratios BCR-ABL1/GUSB of the reference test ranged from −3.52 log10% (0.0003 BCR-ABL1/GUSB%) to 1.36 log10% (23 BCR-ABL1/GUSB%). Among the 28 patient samples 67.8% of the cases (19/28) were shown to be detectable and quantified by both tests, while 10.7% of cases (3/28) were shown to be negative by both tests. 14.3% of cases (4/28) were detected by reference test and undetected by Blink test. 7.1% of cases (2/28) were undetected by reference test and detected by Blink.

A Bland Altman plot analysis was performed using the 19 quantitative results. The Bland Altman plot in Figure 6 shows a small constant bias between both methods across the complete measuring range of 0.06 log10%. This bias can be eliminated with a conversion factor. The figure also shows the upper and lower 2SD of the mean difference that was observed with SD = 0.28 log10%. There is no indication that the variation is dependent on the BCR-ABL1 transcript level.

**Figure 6.**
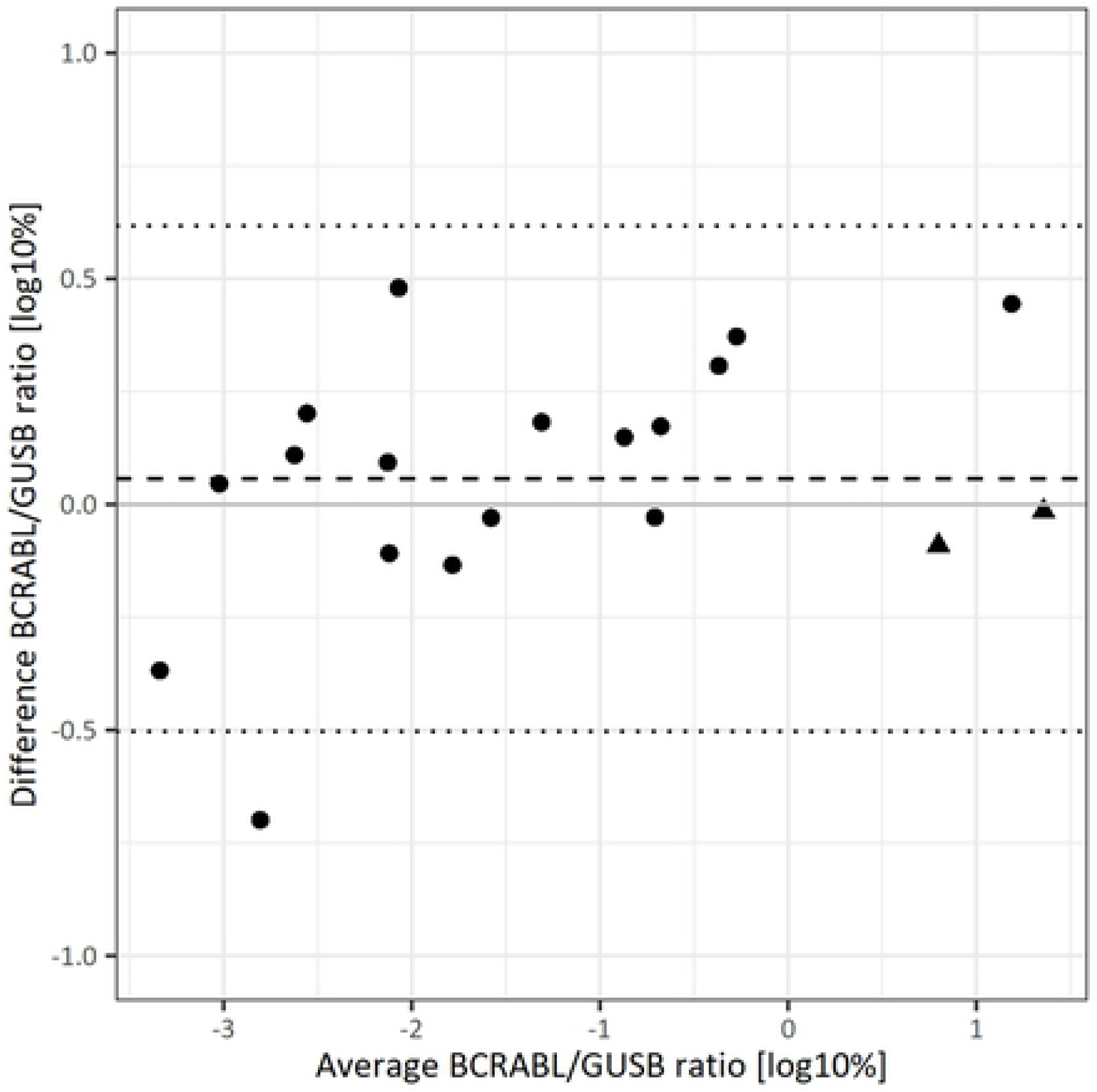
Bland Altman plot for data obtained with the reference method and with the bead nanoreactors. (dots: Poisson analysis of digital PCR data for BCR-ABL1, triangles: Ct-value analysis of quantitative real-time PCR for BCR-ABL1)

Figure 7 shows the Deming regression fit for the same data. The slope was determined with 1.08 with 95% CI from 0.99 to 1.21. The intercept was 0.17 with 95% CI from 0.03 to 0.38. Excellent concordance between the nanoreactor bead assay and the reference with a strong linear relationship (Pearsons’ r = 0.983, p = 5×10^−14^) has been observed.

**Figure 7.**
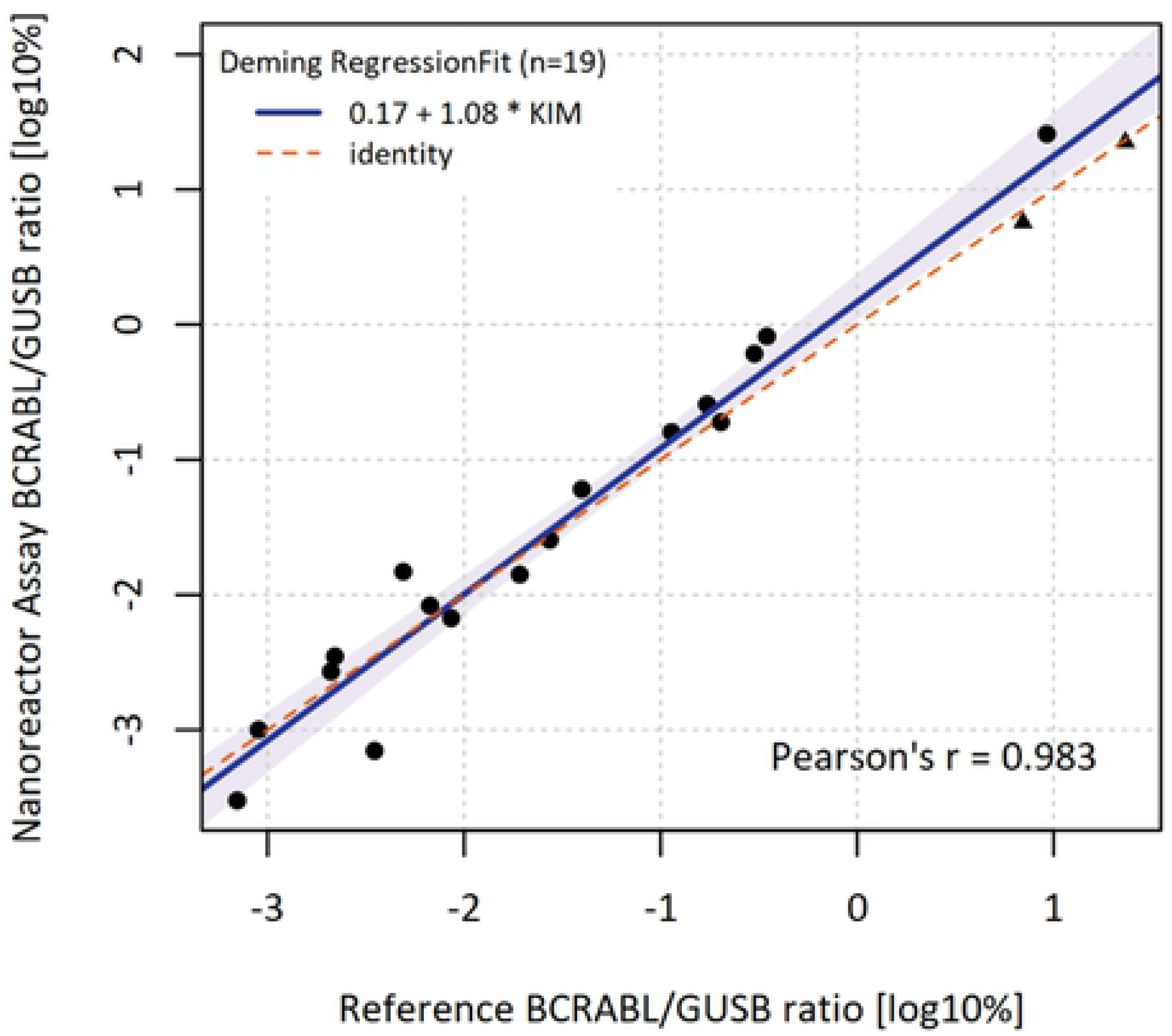
Deming Regression for data obtained with the reference method and with the bead nanoreactors. (dots: Poisson analysis of digital PCR data for BCR-ABL1, triangles: Ct-value analysis of quantitative real-time PCR for BCR-ABL1, Ref = reference)

## Conclusions

Nanoreactor beads are a new class of molecular reagents enabling microfluidics free digital molecular assays. The theoretical upper limit of quantification based on Poisson analysis corresponds to approximately 10x the number of bead-nanoreactors employed in the assay. To overcome this limitation, we supplemented the analysis of digital PCR with real-time quantitative PCR analysis for target concentrations exceeding the upper limit of digital quantification. This approach enabled us to design an assay for the quantification of BCR-ABL1, GUSB, and for the ratios BCR-ABL1/GUSB with a dynamic range of >6 orders of magnitude and to test the assay on clinical peripheral blood samples. The results indicate that the developed approach provides for high sensitivity and, to our knowledge, an unprecedented measurement range in a single reaction. Moreover, the assay is simple to perform and delivers quantitative results comparable with the current laboratory standard.

## Acknowledgements

This work has been supported by the Free State of Thuringia and co-financed with funds of the European Union under the European Regional Development Fund under grant 2017 FE 9005.

We wish to thank the patients who have agreed to provide the samples for our method comparison. For excelent technical work, we like to thank Ines Engelman (BLINK AG, Jena, Germany) and Anja Waldau (Universitätsklinikum Jena, Klinik für Innere Medizin II, Abteilung Hämatologie und Internistische Onkologie, Jena, Germany).

## Suporting information

S1 Table 1

